# Autolab HBH: A Rapid Hyperbaric Heating Device for Streamlined, PCR-Ready Sample Preparation Across Diverse Biological Matrices and Organisms

**DOI:** 10.64898/2026.01.21.700867

**Authors:** Alem Abebe, Brian Miller, Tammo Heeren, Sarkis Babikian, Kirk Allen, Jacob Hambalek, David Wright, Regis Peytavi

## Abstract

Traditional nucleic acid extraction methods are costly, lengthy, and highly variable depending on the complexity of the sample matrix or the organism of interest. Workflows may exceed twenty steps, require separate kits for RNA and DNA, and demand expensive instrumentation, creating barriers to both speed and scalability.

The Autolab^TM^ HBH system addresses these limitations by using hyperbaric heating (HBH) to achieve temperatures above 100 °C in a sealed, pressurized environment through induction heating, enabling rapid lysis of diverse organisms and neutralization of macromolecular PCR inhibitors within minutes. The combination of extreme heat and HBH-optimized lyophilized reagents rapidly inactivates nucleases while preserving free nucleic acids. The workflow is streamlined to two steps: heating up to 1 mL of sample in the proprietary HBH bullet, followed by a brief centrifugation to pellet additives. The resulting supernatant is immediately compatible with real-time reverse transcription polymerase chain reaction (RT-PCR) and other downstream molecular assays.

Here, we evaluate the system’s broad compatibility with diverse sample buffers, matrices, and organisms. Comparative testing was conducted alongside Qiagen extraction methods to benchmark performance.

## Introduction

Accurate detection of infectious organisms increasingly depends on polymerase chain reaction (PCR) based methods, creating a strong need for sample preparation strategies that are rapid, cost-effective, and capable of generating inhibitor-free, PCR-ready lysates. Traditional approaches, such as culture, require several days, while widely used extraction kits (e.g., Qiagen’s DNA and RNA kits) typically require 0.5–3 hours, regardless of sample source or organism type. Workflows using these kits involve multiple handling steps: for example, the RNeasy Mini Kit requires 6–12 steps depending on sample type (1), and the AllPrep PowerFecal Pro DNA/RNA Kit involves 17 steps (2). These protocols are further complicated by the presence of PCR inhibitors commonly found in clinical, veterinary, and environmental samples, necessitating additional purification steps. Moreover, most commercial extraction kits are tailored to specific matrices or organism classes and require separate workflows for RNA and DNA, increasing operational complexity and cost.

The challenges associated with traditional sample preparation were central to the development of hyperbaric heating (HBH), originally pioneered within AMDI’s Fast PCR system (3). The overarching aim of Fast PCR was to create a true sample-to-answer system capable of detecting a broad range of pathogens in diverse sample types in less than 10 minutes, while preserving the core strengths of PCR—namely, analytical sensitivity and specificity. While advances in high-speed thermoresistant polymerases have made fast PCR feasible, sample preparation remained the key bottleneck preventing total assay times from falling below the 10-minute threshold.

Nucleic acid sample preparation must accomplish two essential tasks: disruption of microbial cell walls to release genomic material and mitigation of PCR inhibitors. Existing strategies generally fall into two categories. The first involves simple heat lysis followed by dilution to mitigate inhibitors; although straightforward and automation-friendly, this approach often requires 5–10 minutes of heating near the boiling point and can compromise sensitivity through dilution. The second relies on multistep extraction workflows that deliver high-quality nucleic acids but require harsh chemical reagents, binding resins, wash buffers, and several sequential steps, making them labor-intensive, slow, and challenging to automate.

HBH was developed to bridge these two approaches by enabling rapid and efficient lysis under elevated temperature and pressure. By heating samples in a sealed metallic vessel through induction, HBH reaches temperatures of 100–130 °C within seconds—sufficient to lyse even difficult-to-disrupt organisms. Pressurization allows the sample to reach these temperatures, while induction heating provides fast and homogeneous ramp rates. A proprietary blend of lyophilized chelators and proteins protects nucleic acids, including thermally sensitive RNA, from degradation, while extreme heat and reagent chemistry work together to neutralize macromolecular PCR inhibitors such as proteins and nucleases. What remains is nucleic acid suitable for direct PCR without purification, enabling a rapid workflow compatible with the goals of fast molecular testing. Taken together, the simplicity, efficiency, low hands-on requirements, and speed of HBH underscored its potential as an independent Research Use Only (RUO) sample preparation platform, prompting the development of the Autolab HBH system.

Building on these principles, the Autolab HBH system offers a rapid, versatile, and simplified sample preparation platform capable of producing PCR-ready RNA and DNA in under two minutes. The system supports a wide range of specimen types—including nasal and buccal swabs, vaginal swabs, urine, and stool—processed in diverse buffer formulations such as Tris-EDTA buffer (TE), AMDI Sample Buffer, BD universal viral transport system (UVT), and MicroTest^TM^ M4RT. Preliminary evaluations also indicate compatibility with selected veterinary and environmental matrices (data not presented here). While certain sample types may require minimal preprocessing, the overall workflow remains substantially streamlined compared to traditional extraction methods.

The Autolab HBH instrument provides programmable control of key lysis parameters— temperature (100–150 °C), hold time (0–60 s), and input volume (300–1000 µL)—allowing optimization across diverse organisms, including more resistant microbes such as spores. Samples are loaded into proprietary HBH bullets containing lyophilized reagents and heated under hyperbaric conditions via induction. Following a brief centrifugation step to pellet additives, the supernatant is immediately compatible with downstream molecular assays.

Together, these features address longstanding limitations in traditional sample preparation and align with the original Fast PCR objective: enabling rapid, sensitive, and broadly compatible molecular detection by removing sample prep as the primary bottleneck in ultrafast PCR workflows.

This paper provides an overview of the versatility and overall compatibility of a wide range of sample types using the Autolab HBH system, its effective inactivation of PCR inhibitor nucleases, as well as sample stability post Autolab HBH processing.

## Materials and Methods

### Sample Types and Collection

Buffer compatibility with the Autolab HBH workflow was evaluated using TE, BD Universal Viral Transport (UVT), M4RT, and AMDI Sample Buffer (Table 1).

**Table 1:**
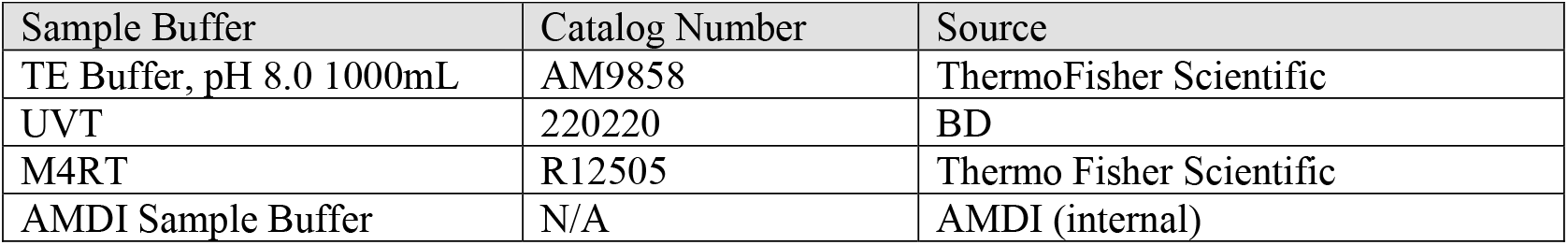
Buffers evaluated, including catalog numbers and suppliers.

A range of specimen types—nasal swab, buccal swab, urine, and stool—were evaluated to assess the versatility of the HBH lysis method. Nasal swab, buccal swab (Catalog No. 25-3406-H, Puritan), and urine samples were collected internally under IRB-approved protocols (IRB ID: 21-AMDI-101). Each nasal and buccal swab was resuspended in 1.5 mL of the designated buffer, aliquoted into single-use volumes, and stored at −20 °C until use. Depending on the study, either single-donor material or pooled samples from 5–10 donors were utilized. Stool samples were sourced from PRECISION for Medicine (Norton, MA) and stored at −80 °C per vendor guidelines.

### Organisms and Contriving of Samples

To evaluate the broad applicability of the HBH lysis method across diverse microbial groups, strains from six species were included in this study. Microorganism stocks were obtained from Microbiologics (San Diego, CA), and ZeptoMetrix (Buffalo, NY). Viral, bacterial, and fungal stocks were quantified by droplet digital PCR (ddPCR) and/or titered in TCID_50_/mL, or CFU/mL by the vendor. Details regarding strain identity and source are provided in Table 2.

**Table 2:** Microbial species evaluated, strains and suppliers.

| Microbial Organism | Strain | Source |
| --- | --- | --- |
| SARS-CoV-2 | USA/WA1/2020 | Microbiologics |
| Human Respiratory Syncytial Virus (RSV) | A2 | Microbiologics |
| Influenza virus B | B/Phuket/3073/2013 | Microbiologics |
| Candida auris | MMX 9869 | Microbiologics |
| Candida albicans | Z006 | ZeptoMetrix |
| Campylobacter jejuni | Z086 | ZeptoMetrix |

Organism stocks were stored at −80 °C, and sample matrices at −20 °C until use. On the day of each study, single-use aliquots of both organisms and sample matrices were thawed. Test samples were contrived by dilution of microorganism stocks into negative matrices at the appropriate concentration, while keeping the organism stocks, negative matrix, and samples on ice.

### Autolab HBH Sample Preparation Workflow

The Autolab HBH K01 kit, designed for 300–600 µL input volumes, was used for all studies. Samples contrived with the designated organisms were aliquoted at 400 µL into HBH bullets and maintained at room temperature prior to processing. Within one hour of preparation, samples were processed on the Autolab HBH instrument using a target temperature of 130 °C and a 0-second hold time. After HBH treatment, samples were briefly centrifuged using a standard benchtop centrifuge, and the resulting supernatant was maintained on ice until all conditions were ready for PCR setup.

### RNase Activity Assay

RNase activity in nasal swab matrix was assessed using the RNaseAlert™ fluorogenic assay (Catalog No. 11-02-01-02, Integrated DNA Technologies, IDT) according to the manufacturer’s recommendations with minor modifications. Pooled nasal swab matrix prepared in AMDI Sample Buffer was aliquoted into HBH bullets containing lyophilized processing reagents and subjected to hyperbaric heating for 0, 5, 10, or 15 seconds. Untreated samples served as matrix controls. Following HBH processing, samples were serially diluted to minimize matrix effects prior to assay input. RNase-positive and RNase-negative controls were prepared using RNase A and nuclease-free water, respectively. Reaction mixtures containing fluorogenic RNA substrate and reaction buffer were incubated with sample dilutions, and fluorescence was measured kinetically over 30 minutes using a microplate reader with fluorescein detection settings. RNase activity was quantified based on fluorescence signal over time.

### Comparison of sample buffer compatibility

Nasal swab specimens collected from a single donor and resuspended in AMDI Sample Buffer, TE, M4RT, or UVT were spiked with 1,000 copies/mL of inactivated SARS-CoV-2. Samples (N=2 per buffer) were processed using the Autolab HBH workflow and analyzed by RT-PCR on the QuantStudio platform.

### Comparator Extraction Workflows (Qiagen)

For comparative evaluation of SARS-CoV-2 detection, samples were extracted using the QIAcube Connect automated platform with the QIAamp Viral RNA Kit (Cat. No. 52904) following the manufacturer’s handbook procedures.

Comparative studies with *Candida albicans* employed the DNeasy Blood and Tissue Kit (Cat. No. 69504) on the QIAcube Connect. Zymolyase 20T (MP Biomedicals™, Cat. No. MP08320921) was prepared at a final concentration of 10 U/mL in the manufacturer-recommended buffer (1000 mM Tris, 500 mM EDTA, 500 mM TCEP). The DNeasy Mini Kit workflow includes a Zymolyase incubation step within the automated protocol. For samples processed using the Autolab HBH, an identical Zymolyase incubation (30 minutes at 37 °C) was performed prior to HBH treatment to ensure methodological equivalence across workflows.

For the *Candida albicans* RNA comparison, samples were processed manually using the RNeasy Mini Kit (Cat. No. 74104, Qiagen, Germany) according to the manufacturer’s protocol. For conditions requiring mechanical disruption, 0.5-mm glass beads (Cat. No. 13116-400, Qiagen, Germany) were used for bead beating on the BeadBug™ Microtube Homogenizer (Cat. No. Z763705, The Lab Depot, GA) prior to either Qiagen or Autolab HBH processing. An additional comparison condition incorporated the QIAshredder (Cat. No. 79654, Qiagen, Germany) for lysate homogenization following the RNeasy handbook instructions.

### PCR and RT-PCR Assays

PCR amplification was performed using the Applied Biosystems™ QuantStudio™ 5 Real-Time PCR System. Reaction mixes contained KAPA3G PCR Buffer (Roche; 1× final, Material No. 09160914103), KAPA3G HotStart DNA Polymerase (Roche; 0.08 U/µL, Material No. 08918651103), MgSO_4_ (Sigma Aldrich; 4.5 mM, Product No. M3409), and dNTPs (Thermo Fisher Scientific; 0.2 mM, Catalog No. R1122). SuperScript™ IV Reverse Transcriptase (Thermo Fisher Scientific; 0.4 U/µL, Catalog No. 18090200) was added for RT-PCR assays.

Forward and reverse primers (1 µM each) and probes (0.45 µM) were sourced from IDT. Primer/probe sequences for SARS-CoV-2, RSV A2, and Influenza B were internally designed, whereas sequences for *C. auris* (4), *C. albicans* (5,6), and *Campylobacter jejuni* (7) were obtained from previously published assays. Published sequences incorporated in this study are provided in Table 3.

**Table 3:**
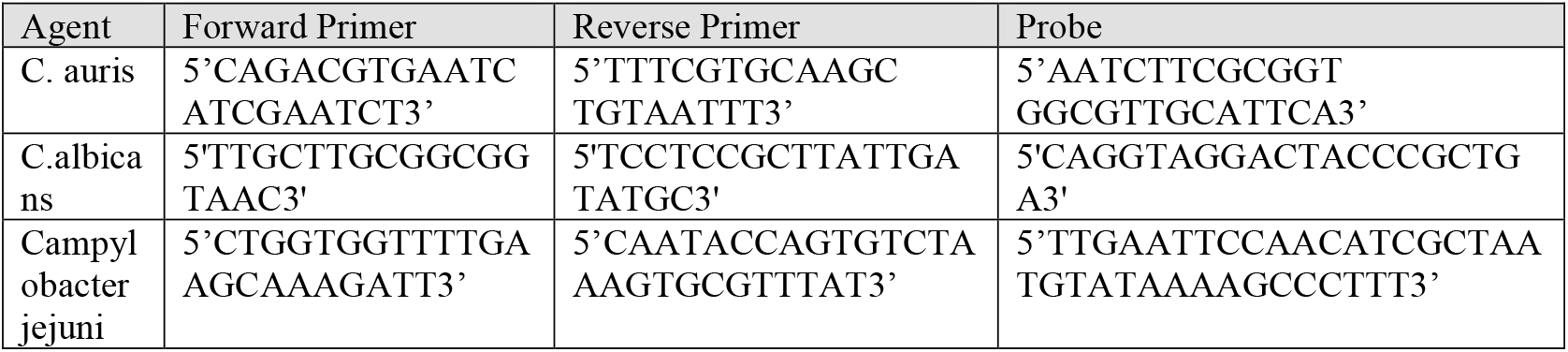
Published primers and probe sequences incorporated in this study.

Total reaction volume was 25 µL, comprising 19 µL of sample and 6 µL of PCR master mix. Ct values reported in this study represent cycle thresholds manually set for comparison across different studies, where lower Ct values indicate higher nucleic acid in the input sample.

## Results

### RNase Activity

RNase activity in pooled nasal swab matrix was evaluated using a fluorogenic RNA probe assay to determine the effect of increasing durations of hyperbaric heating. The RNase-positive control maintained maximal fluorescence throughout the assay period, while the RNase-negative control remained at baseline, confirming assay performance. HBH-treated samples demonstrated a time-dependent reduction in RNase activity, with complete suppression observed following 15 seconds of hyperbaric heating (Figure 2). This indicates that the thermal conditions generated by the Autolab HBH workflow effectively inactivate endogenous RNases present in nasal swab matrix, supporting preservation of RNA integrity for downstream molecular applications.

**Figure 1:**
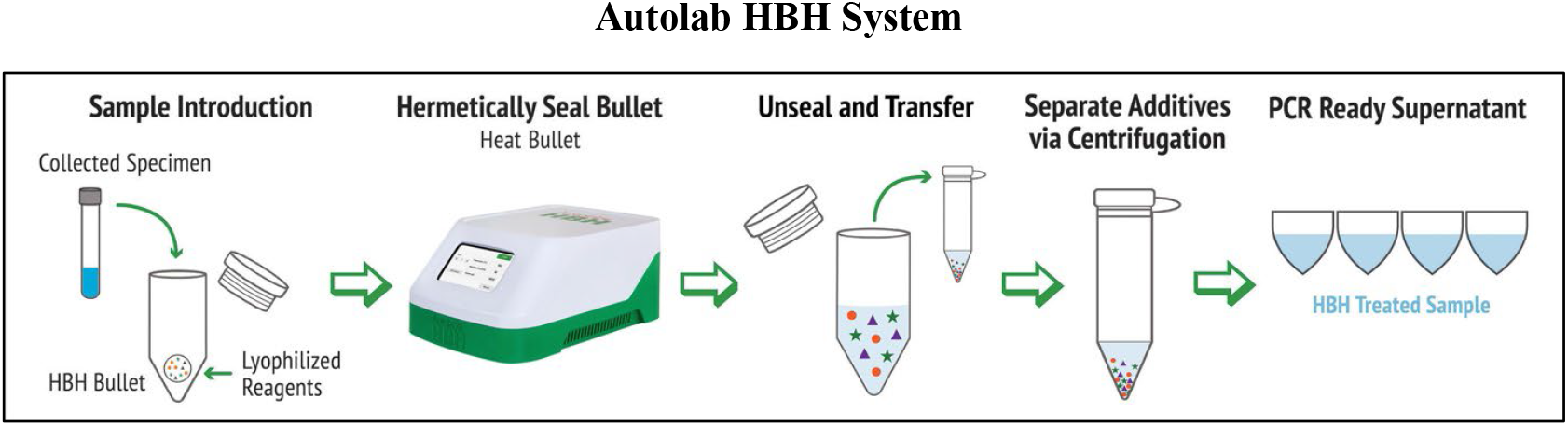
The standard Autolab HBH system workflow consists of two steps: (1) Heat up to 1mL of sample in the HBH bullet and (2) Briefly centrifuge to pellet lyo additives. The supernatant is then ready for RT-PCR, as well as additional molecular testing.

**Figure 2:**
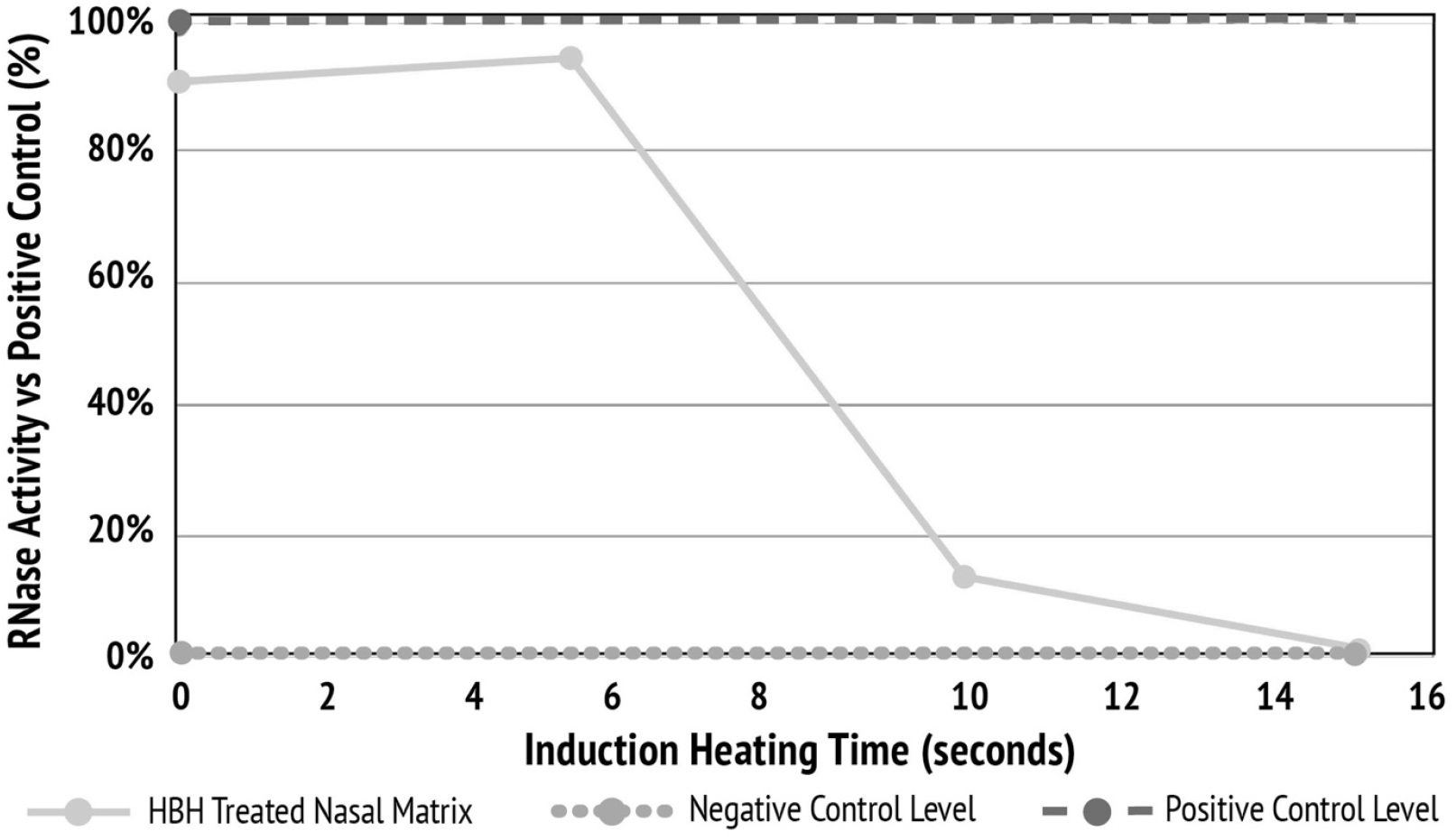
Assessment of RNase activity in nasal swab matrix. RNase activity was quantified by measuring fluorescence following a 10-minute incubation with a fluorescent RNA probe. RNase activity in pooled nasal swab matrix decreased with increasing HBH exposure time, reaching complete inactivation after 15 seconds. RNase-positive and RNase-negative controls are shown for reference.

### Comparison of Sample Buffer Compatibility

To evaluate buffer compatibility with the Autolab HBH workflow, nasal swab specimens were processed across multiple commonly used transport and storage buffers. Ct values differed by no more than two cycles across all conditions, demonstrating consistent assay performance independent of buffer composition (Figure 3).

**Figure 3:**
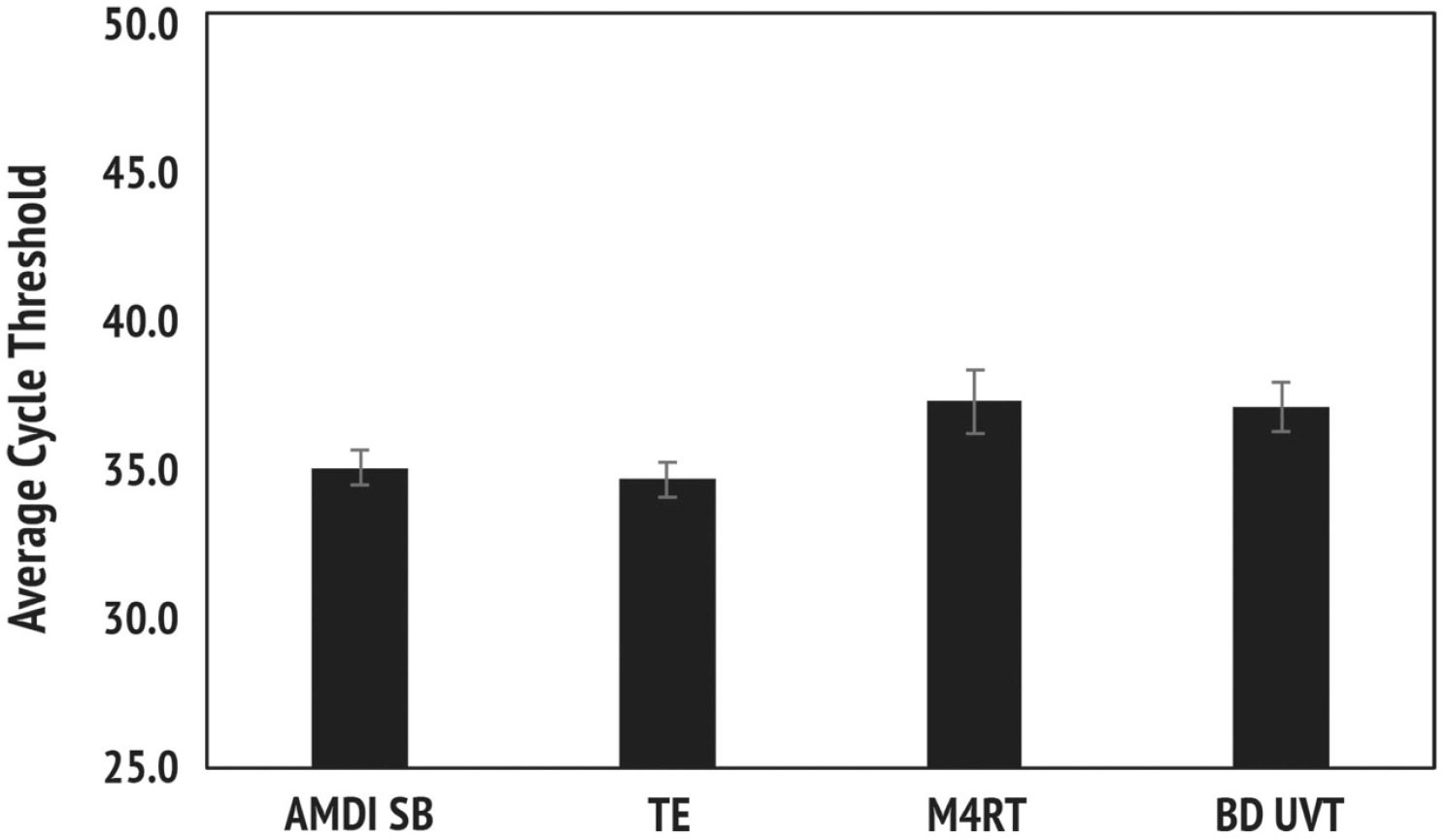
Comparison of sample buffer compatibility with the Autolab HBH. Nasal swab specimens collected from a single donor and resuspended in AMDI Sample Buffer (AMDI SB), TE, M4RT, or UVT were spiked with 1,000 copies/mL of inactivated SARS-CoV-2. Samples (N=2 per buffer) were processed using the Autolab HBH workflow and analyzed by RT-PCR on the QuantStudio platform.

### Comparator Extraction Workflows (Qiagen)

To compare the performance of Autolab HBH with an established automated extraction platform, pooled nasal swab matrix spiked with inactivated SARS-CoV-2 was processed in parallel using Autolab HBH and the Qiagen QIAcube workflow. At higher input concentrations (10,000 and 1,000 copies/mL), both methods produced comparable Ct values (Figure 4). At the lowest input level tested (100 copies/mL), Autolab HBH demonstrated improved sensitivity, yielding earlier Ct values and reduced variability relative to the QIAcube (Figure 4). Consistent with these findings, Autolab HBH achieved 100% detection at all concentrations tested, whereas the QIAcube detected 83% of samples at 100 copies/mL (Table 4).

**Table 4:** SARS-CoV-2 detection rates following Autolab HBH and Qiagen QIAcube processing.

| Viral Load | % Detected Autolab HBH | % Detected QIAcube |
| --- | --- | --- |
| 100 cp/mL | 100% | 83% |
| 1,000 cp/mL | 100% | 100% |
| 10,000 cp/mL | 100% | 100% |

**Figure 4:**
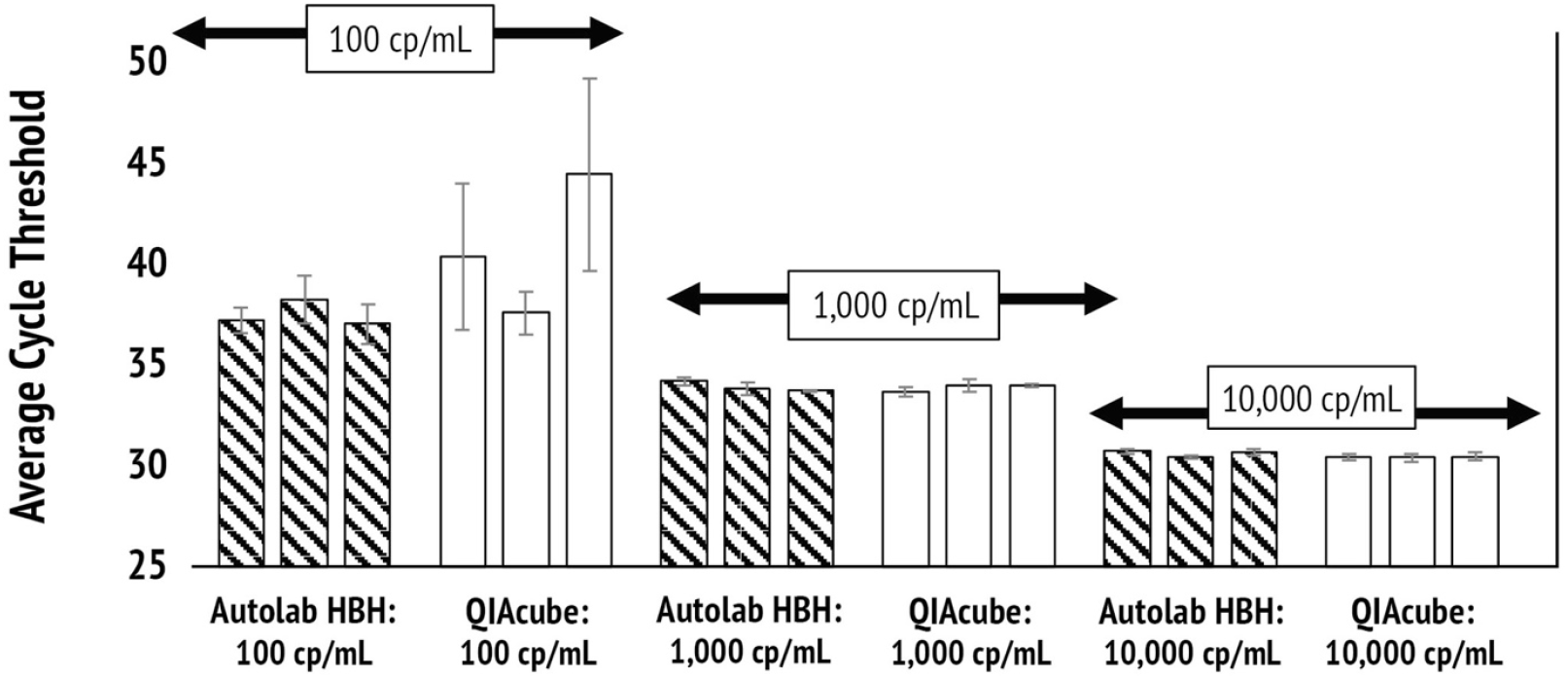
Comparative performance of Autolab HBH and Qiagen’s QIAcube. Pooled nasal swab matrix was spiked with inactivated SARS-CoV-2 at 10,000, 1,000, and 100 copies/mL and processed in parallel using the Autolab HBH workflow and the Qiagen QIAcube extraction system. Processed samples were analyzed by RT-PCR on the QuantStudio platform.

Following comparison of Autolab HBH with automated Qiagen extraction for viral targets, we next evaluated fungal sample preparation performance relative to Qiagen’s recommended workflows using *Candida albicans* in buccal swab matrix. For RNA-based detection, Autolab HBH was compared with the Qiagen RNeasy Mini Kit using both bead-beating and QIAshredder-based homogenization strategies (Figure 5a). At higher input concentrations (10,000 and 1,000 CFU/mL), Autolab HBH—with or without bead beating—produced Ct values comparable to the RNeasy bead-beating workflow, whereas the QIAshredder-based approach resulted in markedly delayed amplification.

**Figure 5a.**
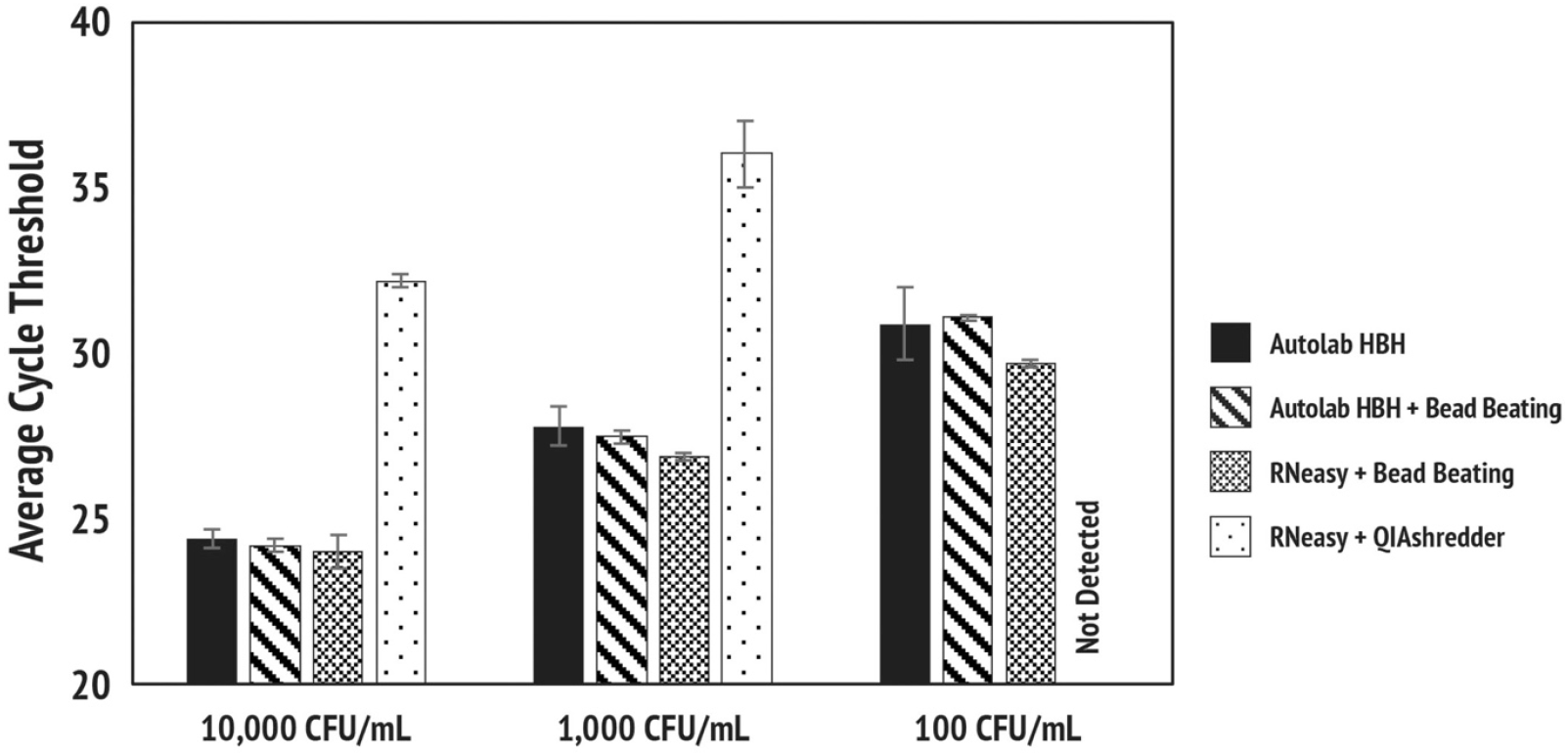
Detection of *Candida albicans* in buccal swab matrix: Autolab HBH versus Qiagen RNeasy. Pooled buccal swab material prepared in TE buffer was serially spiked with *C. albicans* at 10,000, 1,000, and 100 CFU/mL. Samples were processed in parallel using the Autolab HBH workflow with or without bead beating, the Qiagen RNeasy Mini Kit with bead beating, or the RNeasy Mini Kit with QIAshredder-based homogenization. All processed samples were analyzed by PCR on the QuantStudio platform.

A parallel comparison was performed using Qiagen’s DNeasy workflow for DNA purification, which requires enzymatic lysis with Zymolyase (Figure 5b). While the standard Autolab HBH workflow alone yielded slightly delayed Ct values relative to DNeasy, incorporation of a Zymolyase incubation step prior to HBH processing resulted in earlier amplification than all other conditions tested. Together, these results indicate that Autolab HBH provides fungal lysis performance comparable to Qiagen’s recommended workflows and can be effectively integrated with enzymatic pretreatment when enhanced sensitivity is required.

**Figure 5b.**
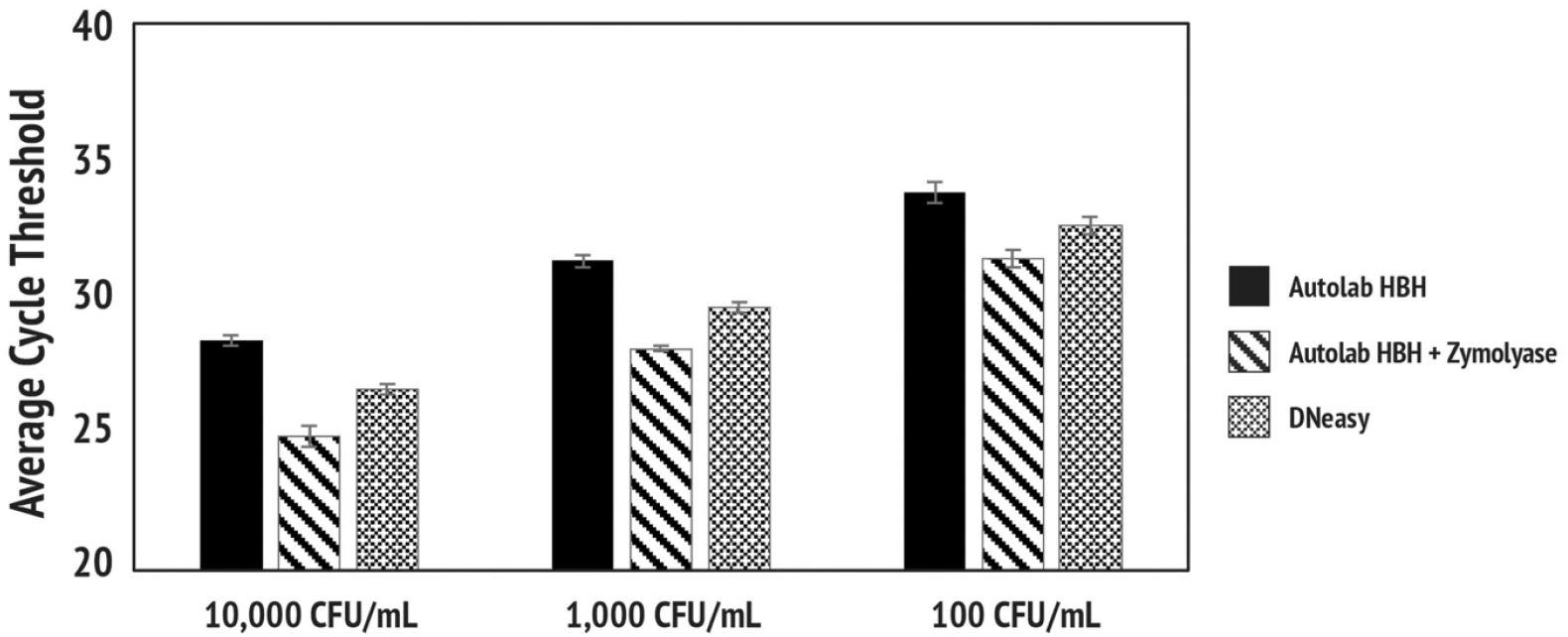
Detection of *Candida albicans* in buccal swab matrix: Autolab HBH versus Qiagen DNeasy. Pooled buccal swab material prepared in TE buffer was spiked with *C. albicans* at 10,000, 1,000, and 100 CFU/mL. Samples were processed using the Autolab HBH workflow with or without prior incubation with Zymolyase or using the Qiagen DNeasy Mini Kit, which incorporates Zymolyase treatment as part of the protocol. PCR analysis was performed on the QuantStudio platform.

### Analytical Sensitivity and Expanded Fungal Target Detection in Buccal Swab Matrix

Following comparison with established extraction workflows, we next assessed the analytical sensitivity of the Autolab HBH for fungal targets in buccal swab matrix. *Candida albicans* was evaluated across serial dilutions spanning more than two orders of magnitude to determine limit-of-detection performance. HBH-processed buccal swab specimens demonstrated consistent PCR amplification across all concentrations tested, including the lowest input level of 25 CFU/mL (Figure 6), indicating robust fungal lysis and target release under standard HBH conditions.

**Figure 6:**
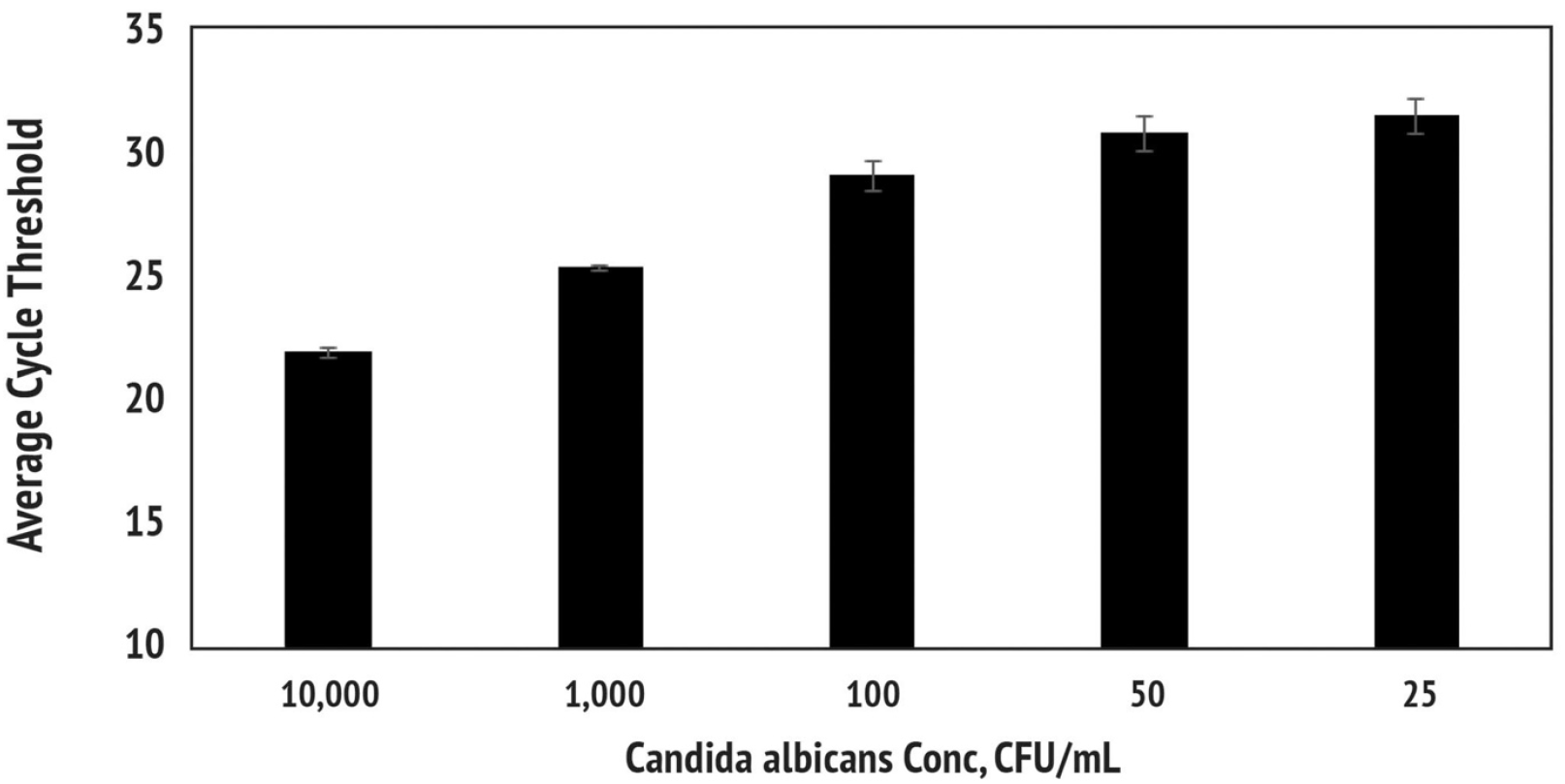
*Candida albicans* detection across a serial dilution in buccal swab matrix. Pooled buccal swab matrix prepared in AMDI Sample Buffer was serially spiked with *C. albicans* from 10,000 CFU/mL down to 25 CFU/mL and processed using the Autolab HBH. PCR analysis was performed on the QuantStudio platform.

To further extend fungal target coverage beyond *C. albicans*, we next evaluated *Candida auris*, an emerging multidrug-resistant pathogen of increasing clinical relevance. Buccal swab specimens spiked with *C. auris* at three input levels yielded reliable PCR detection following HBH processing (Figure 7), confirming effective lysis and compatibility with this additional fungal species. Collectively, these results demonstrate that the Autolab HBH workflow supports sensitive detection of both established and emerging fungal targets in buccal swab matrix without the need for additional purification steps.

**Figure 7.**
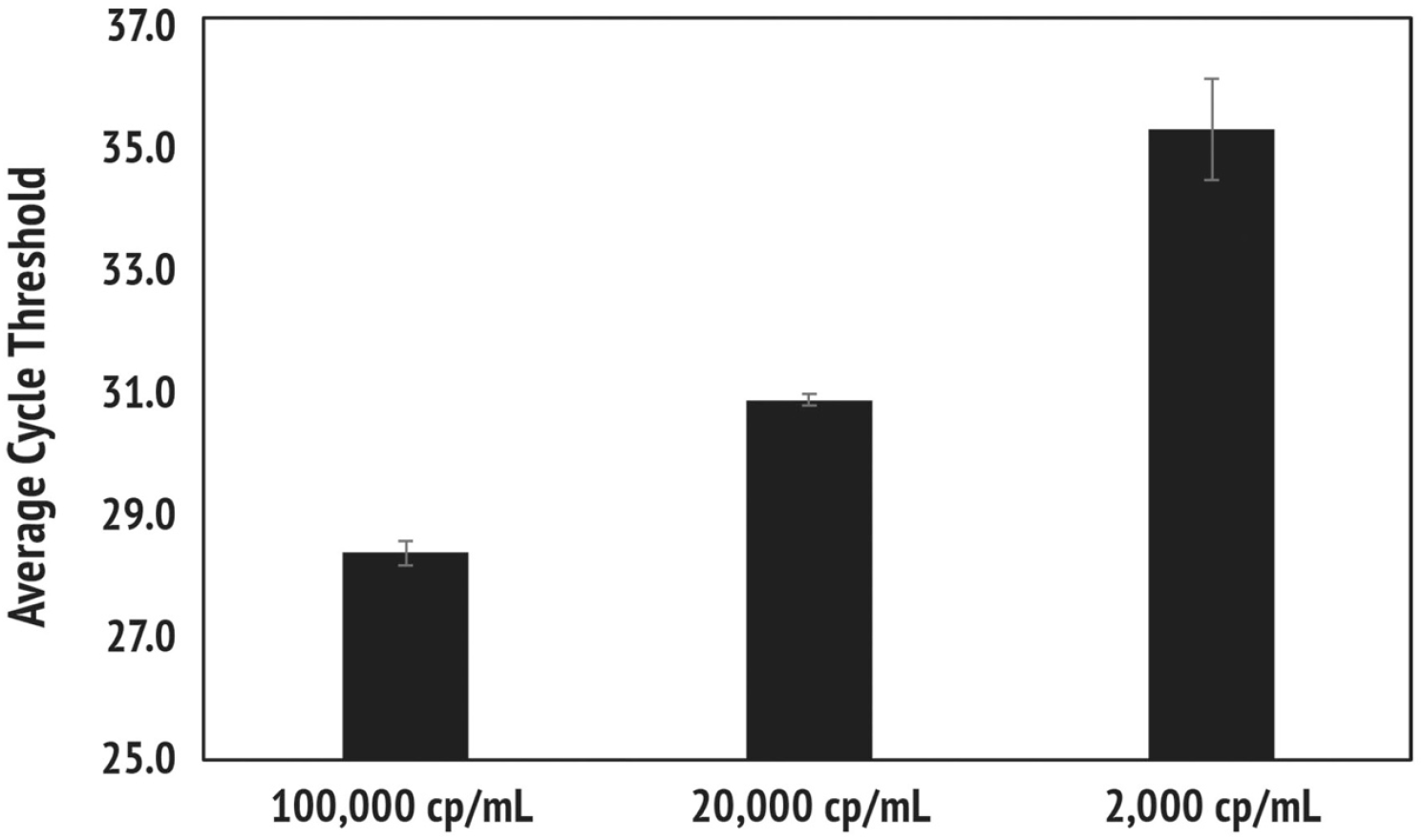
Autolab HBH–mediated detection of *Candida auris* in buccal swab matrix. Pooled buccal swab specimens prepared in AMDI Sample Buffer were spiked with *C. auris* at 100,000 copies/mL, 20,000 copies/mL, and 2,000 Copies/mL and processed using the Autolab HBH workflow. Processed samples were analyzed by PCR on the QuantStudio platform.

### Fungal Target Detection in Urine Matrix

To assess Autolab HBH performance in urine, a matrix known to contain PCR inhibitors, *Candida albicans* detection was evaluated following HBH processing. Urine samples spiked with *C. albicans* were analyzed with and without HBH treatment. Across all tested resuspension volumes, HBH-processed samples yielded substantially earlier amplification compared to untreated controls, demonstrating that thermal processing was necessary to enable reliable PCR detection in this matrix (Figure 8). These findings highlight the ability of the HBH workflow to overcome inhibitory effects associated with urine and support its applicability to non-respiratory clinical specimens.

**Figure 8:**
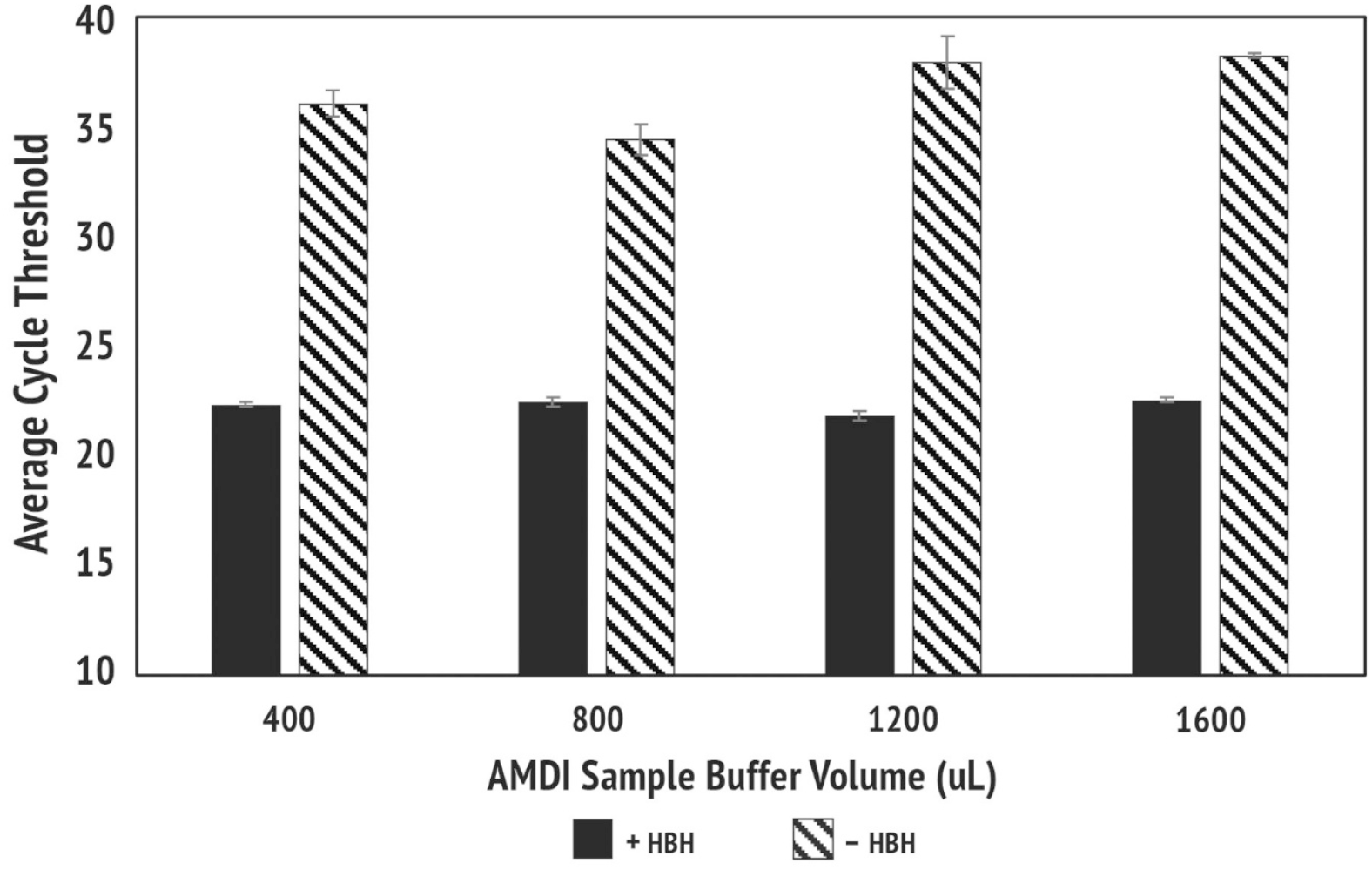
Effect of Autolab HBH processing on *Candida albicans* detection in urine. Urine from a single donor was spiked with *C. albicans* at 10,000 CFU/mL and centrifuged at 3,000 × g for 15 min. Following removal of the supernatant, the pellet was resuspended in 400, 800, 1,200, or 1,600 µL of AMDI Sample Buffer and processed using the Autolab HBH workflow. PCR analysis was performed on the QuantStudio platform using both HBH-treated samples (+HBH) and untreated aliquots (–HBH).

### Detection of Bacterial Targets in Inhibitory Stool Matrices

To further evaluate Autolab HBH performance in highly inhibitory matrices, we assessed detection of the bacterial pathogen *Campylobacter jejuni* in both formed and unformed stool samples. In formed stool, HBH-treated samples and those subjected to conventional heat treatment (100 °C for 10 min) yielded comparable PCR amplification across all tested concentrations, whereas untreated samples failed to amplify (Figure 9a).

**Figure 9a.**
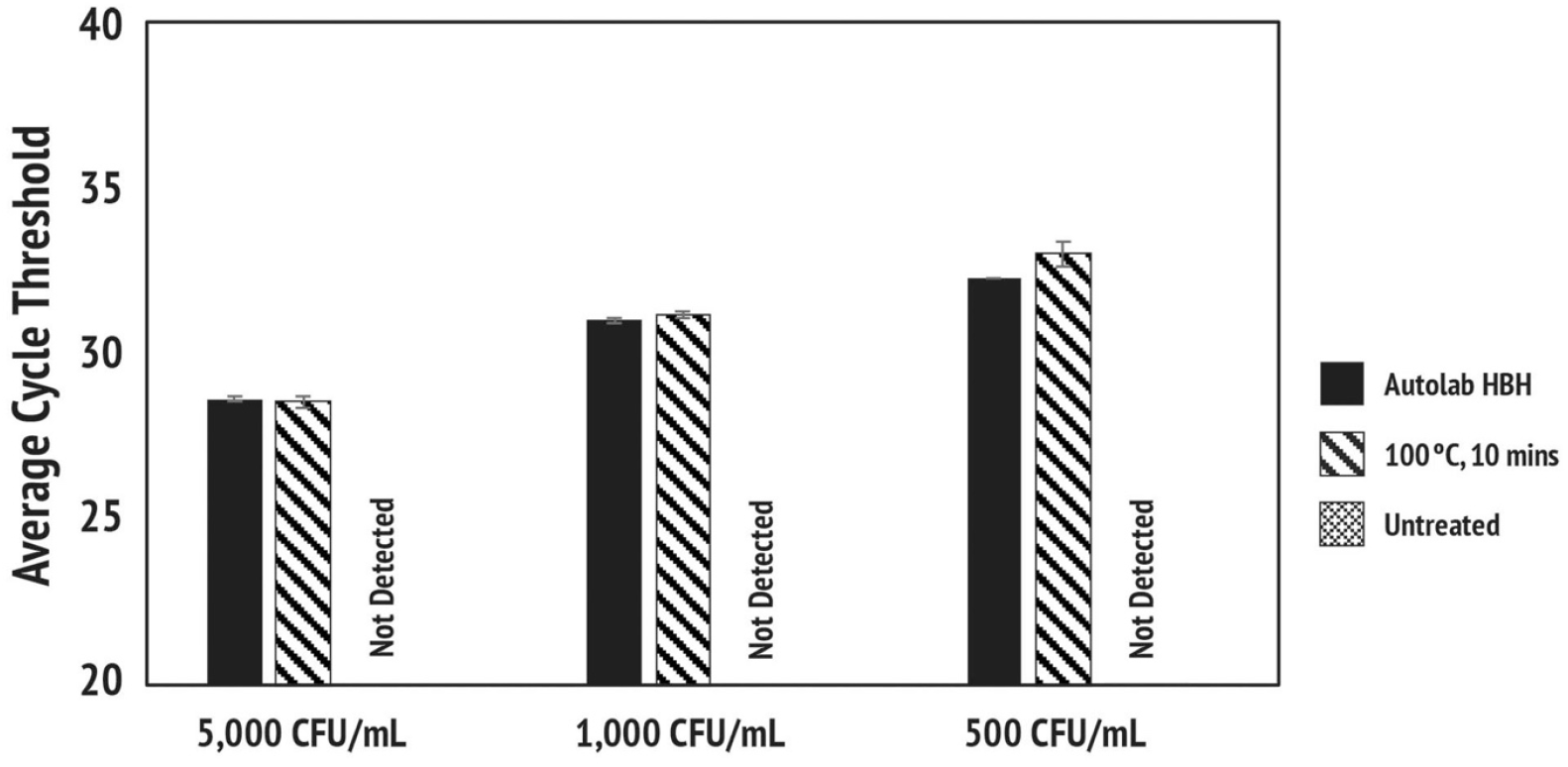
Detection of *Campylobacter jejuni* in formed stool. A swab collected from a single donor’s formed stool was resuspended in 1.5 mL of AMDI Sample Buffer and spiked with *C. jejuni* at 5,000, 1,000, and 500 CFU/mL. Samples were processed in parallel using the Autolab HBH workflow or heating at 100 °C for 10 min, with an untreated aliquot included as a control. PCR analysis was performed on the QuantStudio platform.

In contrast, unformed stool presented a more challenging matrix. While HBH and conventional heat treatment both supported detection at the highest input concentration, only Autolab HBH enabled reliable amplification at lower bacterial loads (1,000 and 500 CFU/mL), with no detectable signal observed for heat-treated or untreated controls (Figure 9b). Together, these results demonstrate that Autolab HBH provides robust bacterial lysis and inhibitor mitigation in stool matrices and offers improved sensitivity relative to standard heat-based processing in more inhibitory sample types.

**Figure 9b.**
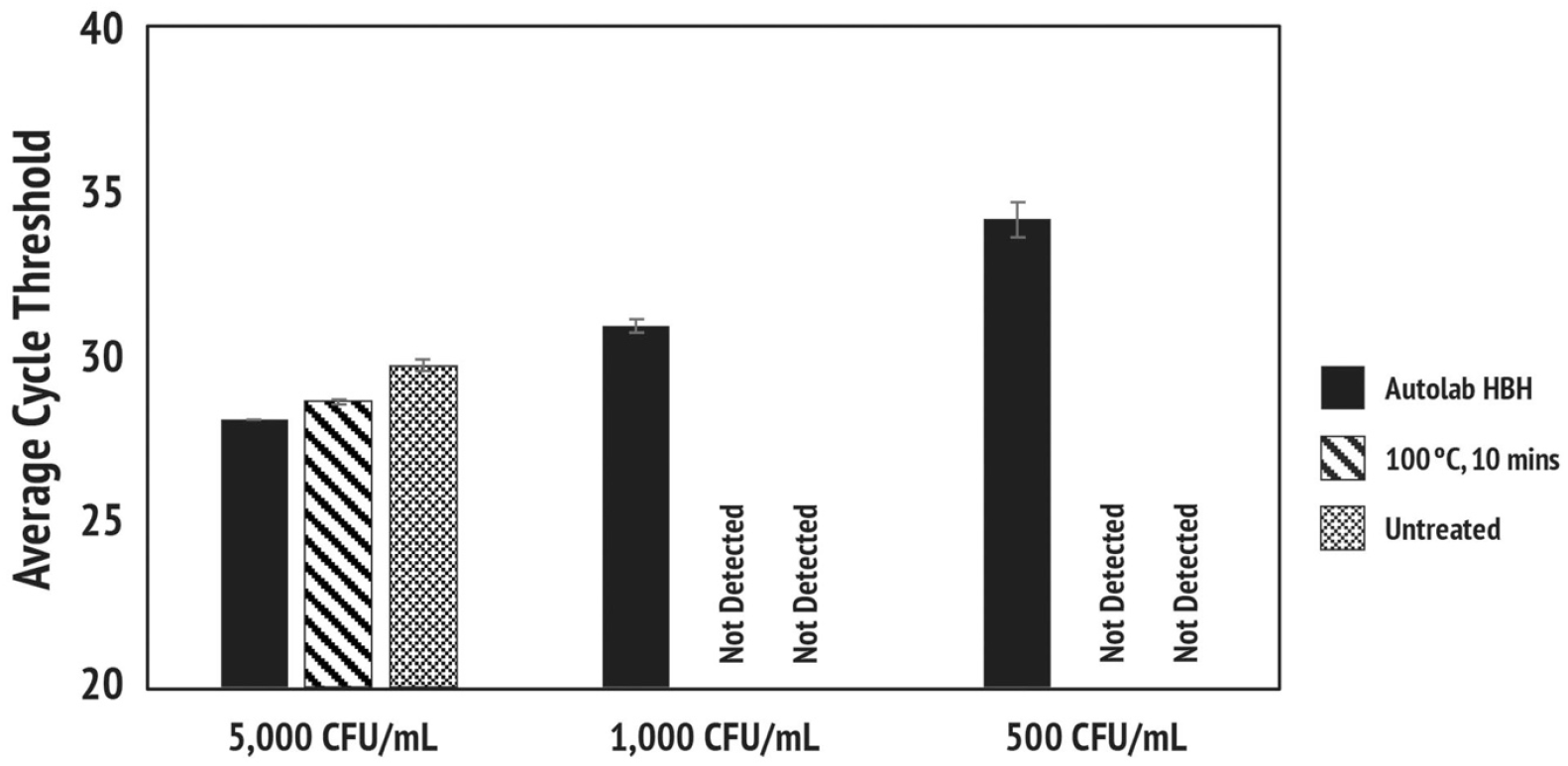
Detection of *Campylobacter jejuni* in unformed stool. Unformed stool samples were processed using the same workflow described for formed stool. Samples were spiked with *C. jejuni* at 5,000, 1,000, and 500 CFU/mL and processed using the Autolab HBH workflow, heating at 100 °C for 10 min, or left untreated prior to PCR analysis on the QuantStudio platform.

## Discussion

Molecular workflows such as PCR, ddPCR, LAMP and all other isothermal nucleic acid amplification depend on effective sample preparation, which conventionally includes cell lysis followed by nucleic acid extraction. Cell lysis can be accomplished using chemical, mechanical, or thermal methods, each aimed at breaching cellular and organelle membranes to release nucleic acids and other intracellular components (8). A variety of methods—chemical, mechanical, and thermal—are used to achieve this disruption. Chemical lysis is widely implemented and typically relies on detergents, chaotrope agents, or enzymes (8). Despite its broad utility, chemical lysis introduces the risk of reagent carryover, and even minimal residual levels (1–10%) can compromise or completely inhibit subsequent amplification reactions (8). In a typical chemical-lysis–based nucleic acid extraction workflow, five to seven core steps (lysis, debris removal, binding, washing, elution) are required, but when including pre-lysis processing and kit-specific variations, the total number of handling steps commonly becomes six to twelve or more.

The Autolab HBH system employs induction-based thermal lysis, allowing samples to reach the high temperatures needed for efficient cell disruption in approximately 15 seconds. The sealed HBH bullet permits temperatures well above 100 °C, enhancing the effectiveness of thermal lysis across diverse organisms. Users may program a 0–60 second hold period at target temperatures between 100 and 150 °C to ensure complete lysis, especially for more resilient microbes such as spores. The lyophilized reagents pre-packaged within the HBH bullets are formulated to stabilize and protect the nucleic acids during and immediately after lysis. Although the standard Autolab HBH workflow does not incorporate a purification step, the data presented here demonstrate that the resulting supernatant remains compatible with PCR or RT-PCR, even at lower organism concentrations.

A key challenge in RNA-based diagnostics is the presence of ribonucleases (RNases), which are abundant in respiratory specimens such as nasal swabs, saliva, and mucus. Active RNases can rapidly degrade free RNA following cell lysis, compromising assay sensitivity. Our results demonstrate that the thermal conditions generated by HBH effectively inactivate endogenous RNases in nasal swab matrix in a time-dependent manner, achieving complete suppression following brief exposure (Figure 2). These findings align with prior reports indicating that rapid, high-temperature processing can mitigate RNase activity and support RNA preservation in complex biological samples (9–11).

To enable broad applicability, we developed AMDI Sample Buffer as a specialized formulation optimized for compatibility with the HBH workflow. Importantly, HBH performance was not limited to this buffer alone; commonly used commercial transport and storage media, including TE, M4RT, and UTM, yielded comparable amplification outcomes (Figure 3). This buffer tolerance highlights the flexibility of the HBH system and supports its integration into existing laboratory workflows without requiring exclusive reagent adoption.

Qiagen extraction kits remain among the most widely used reference methods for laboratory-based nucleic acid purification (12) and served as appropriate comparators in this study. While automated platforms such as the QIAcube Connect reduce some manual handling, they still involve multistep preprocessing and extended run times, contributing to higher cost and operational complexity (12). In comparative analyses using SARS-CoV-2–spiked nasal swab samples, HBH demonstrated analytical performance comparable to, and in some cases exceeding, that of automated extraction while substantially reducing total processing time (Figure 4). These findings suggest that rapid thermal lysis can serve as an effective alternative to traditional extraction workflows for viral targets.

Fungal organisms present additional challenges for molecular detection due to their rigid and compositionally complex cell walls. In comparative studies using *Candida albicans*, a 2-minute Autolab HBH workflow demonstrated lysis performance comparable to Qiagen’s bead-beating-based extraction workflow, which requires approximately 90 minutes of total processing time, while alternative homogenization approaches showed reduced sensitivity (Figure 5a). The enhanced performance observed with enzymatic pretreatment further indicates that HBH can be readily combined with established lysis strategies to improve sensitivity when targeting particularly resilient organisms (Figure 5b). Importantly, HBH alone supported sensitive detection of *C. albicans* across a broad concentration range, underscoring its utility as a stand-alone lysis method for fungal targets (Figure 6).

The applicability of the HBH workflow was further extended to *Candida auris*, an emerging multidrug-resistant fungal pathogen of growing clinical concern (13). Although buccal swabs are not the primary diagnostic specimen for *C. auris*, their use in this study provided a controlled mucosal matrix for evaluating organism-level compatibility and lysis performance. Successful detection across multiple input levels confirms that HBH is effective against this challenging fungal species and supports its broader use for fungal research applications (Figure 7).

Beyond swab-based matrices, HBH demonstrated improved detection sensitivity in more inhibitory sample types. In urine, where PCR inhibition is well documented (14), HBH treatment markedly enhanced amplification compared to untreated samples, even across varying resuspension volumes (Figure 8). Similarly, in stool—one of the most inhibitory matrices for molecular assays due to complex polysaccharides and bile salts (15)—HBH provided robust lysis and improved detection sensitivity, particularly in unformed stool where conventional heat treatment was insufficient at lower target concentrations (Figure 9b). These results highlight the advantage of rapid, high-temperature lysis under pressurized conditions for overcoming matrix-associated inhibition.

## Conclusions

The findings presented here establish the Autolab HBH system as a rapid, low–hands-on-time, versatile, and affordable RUO sample-preparation technology capable of generating PCR-ready lysates across a wide range of organisms, matrices, and buffer conditions—including inhibitory sample types. While the HBH workflow does not purify or concentrate nucleic acids, it effectively replaces the lysis phase of conventional extraction protocols and can be positioned upstream of standard purification workflows when needed. Taken together, the system’s speed, matrix tolerance, and operational simplicity highlight its value both as a stand-alone lysis tool and as a complementary component of broader molecular workflows. Collectively, these attributes position Autolab HBH as a practical and cost-effective RUO solution for research laboratories seeking streamlined sample-preparation strategies.

## References

1. QIAGEN. RNeasy Mini Handbook. QIAGEN, 2023, Germany.

2. QIAGEN. AllPrep PowerFecal ProDNA/RNA Handbook. Qiagen, 2022, Germany.

3. Srinivasan, Aravind, et al. “Multicenter Clinical Comparison of the 10 Minute AMDI™ Fast PCR Mini Respiratory Panel and the Cepheid Xpert Xpress CoV-2/Flu/RSV Plus.” medRxiv, 4 Nov. 2025, 10.1101/2025.11.04.25339522.

4. Centers for Disease Control and Prevention. Real-Time PCR for Identification of Candida auris. CDC, 7 May 2024, https://www.cdc.gov/candida-auris/hcp/laboratories/real-time-pcr-identification.html. (primers and probes, C.auris)

5. Liotti, Flora Marzia et al. “Development of a Multiplex PCR Platform for the Rapid Detection of Bacteria, Antibiotic Resistance, and Candida in Human Blood Samples.” Frontiers in cellular and infection microbiology vol. 9 389. 13 Nov. 2019, doi:10.3389/fcimb.2019.00389 (primers and probes, C.albicans)

6. Wahyuningsih, R et al. “Simple and rapid detection of Candida albicans DNA in serum by PCR for diagnosis of invasive candidiasis.” Journal of clinical microbiology vol. 38, 8 (2000): 3016–21. doi:10.1128/JCM.38.8.3016-3021.2000 (primers and probes, C.albicans)

7. Best, Emma L et al. “Applicability of a rapid duplex real-time PCR assay for speciation of Campylobacter jejuni and Campylobacter coli directly from culture plates.” FEMS microbiology letters vol. 229, 2 (2003): 237–41. doi:10.1016/S0378-1097(03)00845-0 (primers and probes for Campylobacter jejuni)

8. Lee, Soo Min et al. “Chemical Trends in Sample Preparation for Nucleic Acid Amplification Testing (NAAT): A Review.” Biosensors vol. 13, 11 980. 10 Nov. 2023, doi:10.3390/bios13110980

9. Bender, Andrew T et al. “Enzymatic and Chemical-Based Methods to Inactivate Endogenous Blood Ribonucleases for Nucleic Acid Diagnostics.” The Journal of molecular diagnostics: JMD vol. 22, 8 (2020): 1030–1040. doi:10.1016/j.jmoldx.2020.04.211

10. Azmi, Iqbal et al. “A Saliva-Based RNA Extraction-Free Workflow Integrated With Cas13a for SARS-CoV-2 Detection.” Frontiers in cellular and infection microbiology vol. 11 632646. 16 Mar. 2021, doi:10.3389/fcimb.2021.632646

11. Becskei, Attila, and Sayanur Rahaman. “The life and death of RNA across temperatures.” Computational and structural biotechnology journal vol. 20 4325–4336. 8 Aug. 2022, doi:10.1016/j.csbj.2022.08.008

12. Sharma, Pankaj et al. “Experience of quantity and quality of DNA and RNA extraction from limited pediatric blood samples: A comparative analysis of automated and manual kit-based method.” Indian journal of pathology & microbiology vol. 65, 1 (2022): 105–110. doi:10.4103/IJPM.IJPM_946_20

13. Bhargava, Ashish et al. “Candida auris: A Continuing Threat.” Microorganisms vol. 13, 3 652. 13 Mar. 2025, doi:10.3390/microorganisms13030652

14. Khan, G et al. “Inhibitory effects of urine on the polymerase chain reaction for cytomegalovirus DNA.” Journal of clinical pathology vol. 44, 5 (1991): 360–5. doi:10.1136/jcp.44.5.360

15. Schrader, C., A. Schielke, L. Ellerbroek, and R. Johne. “PCR Inhibitors – Occurrence, Properties and Removal.” Journal of Applied Microbiology, vol. 113, no. 5, 2012, pp. 1014–1026.

16. Al-Soud, W A, and P Rådström. “Purification and characterization of PCR-inhibitory components in blood cells.” Journal of clinical microbiology vol. 39, 2 (2001): 485–93. doi:10.1128/JCM.39.2.485-493.2001

17. Hambalek, J. A. Generalizable 15-Second Nucleic Acid Sample Preparation Method via Hyperbaric Heating. Poster 4534, American Society for Microbiology, 18 June 2023, Houston, Texas.

18. Mackay, IM. “Real-time PCR in the microbiology laboratory.” Clinical microbiology and infection: the official publication of the European Society of Clinical Microbiology and Infectious Diseases vol. 10,3 (2004): 190–212. doi:10.1111/j.1198-743x.2004.00722.x

19. Newman, A., and C. Johnson. Candida albicans. StatPearls Publishing, 2022.

